# Super-resolution imaging reveals structurally distinct periodic patterns of chromatin along pachytene chromosomes

**DOI:** 10.1101/023432

**Authors:** Kirti Prakash, David Fournier, Stefan Redl, Gerrit Best, Máté Borsos, Rene Ketting, KiKuë Tachibana-Konwalski, Christoph Cremer, Udo Birk

## Abstract

During meiosis, homologous chromosomes associate to form a unique structure called synaptonemal complex (SC) whose disruption leads to infertility. Information about the epigenetic features of chromatin within this structure at the level of super-resolution microscopy is largely lacking. We combined single molecule localization microscopy with quantitative analytical methods to describe the epigenetic landscape of meiotic chromosomes at the pachytene stage in mouse oocytes. DNA is found to be non-randomly distributed along the length of the SC in condensed clusters. Periodic clusters of repressive chromatin (trimethylation of histone H3 at lysine 27, H3K27me3) are found at 500 nm intervals along the SC, while one of the ends of SC displays a large and dense cluster of centromeric histone mark (trimethylation of histone H3 at lysine 9, H3K9me3). Chromatin associated with active transcription (trimethylation of histone H3 at lysine 4, H3K4me3) is arranged in a radial hair-like loop pattern emerging laterally from the SC. These loops seem to be punctuated with small clusters of H3K4me3 mark with an average spread larger than their spacing. Our findings indicate that the nanoscale structure of the pachytene chromosomes is constrained by periodic patterns of chromatin marks, whose function in recombination and higher-order genome organisation is yet to be elucidated.

## 1 Introduction

Meiosis is a specialized cell division essential for the life cycle of sexual organisms and is needed to generate genetic diversity of haploid gametes. This diversity comes from two major chromosomal events, homologous recombination and chromosome segregation. Recombination initiates by programmed double-strand breaks produced by Spo11 endonuclease in leptotene stage of prophase I.^1^ Homologous chromosomes search for each other to repair the breaks and coalign or pair through this process. The association of homologous chromosomes is later strengthened during zygotene by formation of the synaptonemal complex (SC). The assembly of the SC promotes the close association or synapsis of homologous chromosomes during zygotene. Completion of the SC defines pachytene, when homolog axes are linked along their lengths and reciprocal exchanges of homologous chromosomes result in crossovers, generating genetic diversity. Computational analysis has shown that sites of recombination can be predicted by studying DNA sequence. However, getting the detailed structure of the meiotic chromosomes is essential to understand the mechanistic steps that have to take place in order recombination to happen.^2–4^ While the major recombinase proteins involved in the process of recombination have been localized,^5–7^ the way the structure is constrained by epigenetics is unclear.

In interphase, epigenetics plays an important role in regulating gene expression by modifying chromatin accessibility and recruiting factors to allow genes to become active or silenced.^8^ This is done by regulation of DNA binding proteins called histones. The amino acid tails of histones (H2A, H2B, H3, H4) are susceptible to various post-translational modifications, including acetylation and methylation, which can regulate the compaction of chromatin and modulate gene expression. Post-translational modifications of histones have been shown to contribute to gene expression and recombination during meiosis, but how the structure of the meiotic chromosome is regulated by them is still elusive. A major regulator of meiotic recombination is the H3K4 methyltransferase PRDM9.^9^ The presence of H3K4me3 (trimethylation of histone H3 at lysine 4) is associated with recombination, along with H3K4me2 and H3K9ac.^10, 11^ Conversely, other histone marks are depleted at recombination sites, such as H3K27me3 (trimethylation of histone H3 at lysine 27) and H3K9me3 (trimethylation of histone H3 at lysine 9),^11^ leaving one with the expectation that histone modifications might mark different localizations on the meiotic chromosomes. Therefore, we decided to localize DNA, H3K4me3, H3K9me3, H3K27me3 along the length of SC in order to describe the epigenetic status of the meiotic chromosomes at the pachytene stage.

Conventional light microscopy (LM) has shown the colocalization of H3K27me3 and H3K9me3 with SYCP3^12^ but due to the resolution limitations of LM (∼ 200 nm), the precise distribution of these marks along the pachytene chromosome has not been described yet. With the advent of various super-resolution methods,^13^ in particular single molecule localisation microscopy (SMLM) based methods^14–16^ and recently developed strategies for labeling DNA,^17, 18^ it is now possible to study DNA structure at the molecular scale. The underlying principle of most SMLM based methods is to label molecules of interest with fluorescent moeities that can reversibly switch between fluorescent state and a stable dark state. This process of switching between states is called ‘blinking’, and allows for optical isolation of single molecule. Since only a fraction of molecules will actually fluoresce at a given time, their precise location can be determined and a reconstructed super-resolved image can be produced. In this work, we use a flavor of SMLM called Spectral Precision Determination Microscopy (SPDM)^19, 20^ that allows for the usage of conventional fluorophores, which enabled us to investigate chromatin organization via localizing DNA and particular histone modifications along the SC.

In order to describe the general shape of chromatin during meiosis, we applied a recently developed method based on UV-induced photoconversion of standard DNA dyes^17^ that generates high quality density maps of chromatin by promoting blinking. Using this strategy in a conjunction with fluorescently tagged antibodies targeted to particular histone modifications, we studied localization patterns of chromatin and epigenetic modification in pachytene spreads. Chromatin was found to display a characteristic clustering pattern of 550-700 nm along the SC. We then investigated the localization of H3K4me3, H3K27me3 and H3K9me3 along the pachytene chromosomes. Interestingly, we found that the histone marks form highly identifiable clusters, with defined localization and periodicity. Large clusters of H3K27-trimethylated chromatin are periodically detected at intervals of ∼ 500 nm along the synaptonemal complex, while one of the ends of the SC, presumably the telocentric end, displays a large and dense cluster of centromeric histone mark H3K9me3. Chromatin associated with potential recombination regions marked by H3K4me3,^11^ is arranged both axially and in radial loop-like patterns along the SC. The arrangement of H3K4me3 along these patterns seem to be discontinuous forming small clusters, similar to the bead-on-the-string model. The average spacing of 200 nm suggests that regions from which these patterns emanate could alternate with H3K27me3 regions. Overall, these findings help to describe the structure of the pachytene chromosomes, which is punctuated by clusters of epigenetically regulated chromatin that may alternatively promote or suppress recombination.

## 2 Results

### Super-resolution imaging of synaptonemal complex proteins

In order to study the chromatin distribution along the meiotic chromosomes in high resolution, we first characterized the structure of the SC that acts as a scaffold for chromatin during pachytene. This served as our benchmark for the subsequent analyses. We localized the distribution of two important components of the SC, namely SYCP1 (the central element of SC) and SYCP3 (the lateral element of the SC), which are critical for the synapsis of homologous chromosomes. For quantitative analysis, we labelled SYCP3 and SYCP1 C-terminus with Alexa Fluor 555, and detected on average 2500 photons per cycle. Localization maps of SYCP3 and SYCP1 proteins were generated by integrating roughly 20,000 observations, each of which captured photons emitted during 100 ms of camera integration time. These observations localized individual fluorophores with an average precision of 11 nm and 16 nm and a Fourier Ring Correlation (FRC) resolution^21^ of 43 nm and 56 nm, for SYCP3 and SYCP1 C-terminus, respectively. For visualisation, we used the mean distance to 20 nearest neighbour molecules to blur individual molecules of SYCPs. For SYCP3, the nearest neighbour distance was 14 nm *±* 5 nm and for SYCP1, this distance was 19 *±* 7 nm (Fig. 1E). Comparison with localisation precision based Gaussian blurring and triangulation is shown in Sup. Fig 1 and Fig. 4D1-D3.

**Figure 1.**
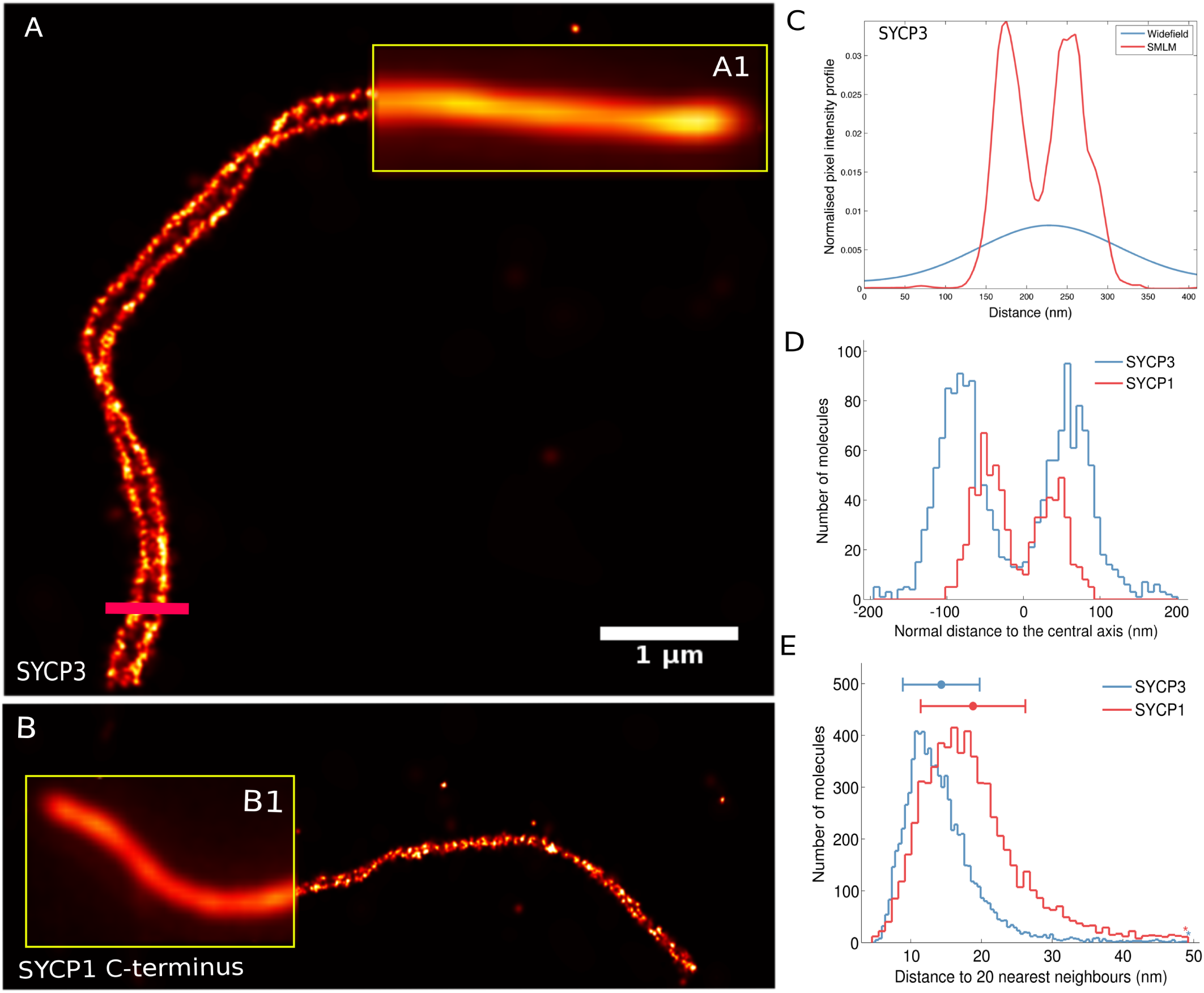
Super-resolution imaging of Synaptonemal Complex Proteins (SYCPs). Single molecule localisation images of SYCP3 (A) and SYCP1 C terminus labelled with Alexa 555 (B) with insets (A1 and B1) showing the widefield equivalent of the underlying localisation image. (C) Plot profile of the transversal distribution of SYCP3, in wide-field imaging (blue plot) and in localization microscopy (red). (D) Plot profile of the bimodal distribution of proteins SYCP1 C-terminal (red plot) and SYCP3 (blue plot). We estimate the inter-strand distance of SYCP3 to be around 180.7 nm and 87.9 nm for the C-terminus of SYCP1 protein. (E) The mean distance to 20 nearest neighbour molecules was used to blur individual molecules of SYCPs. For SYCP3, the nearest neighbour distance was 14.31 nm +/- 5.43 nm and for SYCP1, this distance was 18.81 +/- 7.40 nm.

The images obtained (Fig. 1A-B) show a clear separation of the two lateral elements of the SC and highlight the spatial distribution of individual SYCP3 and SYCP1 proteins along the two strands. Comparison of SMLM image with wide-field image shows that the separation between the two strands of SYCP3 are not resolvable with wide-field microscopy (Fig. 1C). By computing the distance from the maxima of each individual strand of SYCP3, we estimated the distance between the two strands of SYCP3 to be around 181 nm and the width of an individual strand to be 60 nm. Regarding the C-terminus of SYCP1 protein, the distance between the strands was 88 nm and the width 47 nm. Our results are in good argeement with recently reported results, where the width of SYCP3 was reported to be around 56 nm and the distance between strands to be 165 nm.^22^ The authors found the width of SYCP1 C terminus to be 45 nm with overall width of central region (composed of transversal filaments (TFs) and the central element (CE)) to be around 148 nm, which is in line with our observations.

### Non-random distribution of condensed chromatin structures along the meiotic chromosomes

We next analysed, at enhanced resolution, the distribution of chromatin along the pachytene spreads. We immunostained pachytene spreads with anti-SYCP3 (Alexa Fluor 555) and then stained chromatin with Vybrant DyeCycle Violet^18^ (see Materials and Methods for details). Pictures obtained reveal highly condensed chromatin following the SC, with many radial projections (Fig. 2A-C). The highest density of chromatin is found around the central axis of the SC, while chromatin distribution can expand as far as 500 nm away from the center of the SC, though with decreasing density (Fig. 2D).

**Figure 2.**
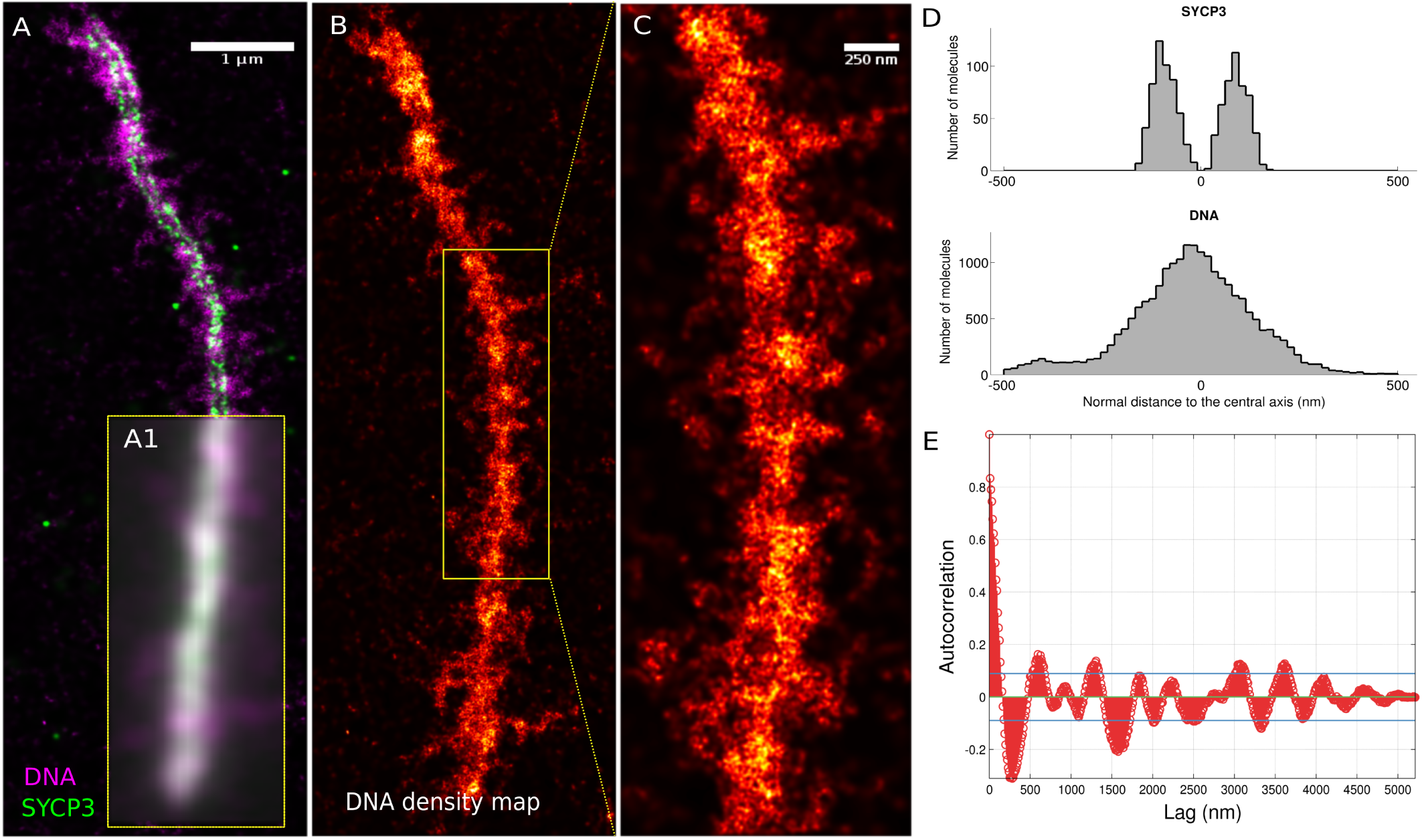
Super resolution microscopy reveals high order clusters of chromatin patterns along the meiotic chromosome. Pachytene spreads were immunostained with anti-SYCP3 (Alexa Fluor 555) and then stained with Vybrant DyeCycle Violet. Two colour SMLM was performed (A), and for comparison, inset (A1) showing the wide-field counterpart of the underlying SMLM image. Diffferential chromatin clusters are highlighted in B and C. (D) In the representative image of meiotic chromosomes stained with Vybrant Dyecycle Violet stained, DNA is normally distributed around the two strands of SYCP3 proteins with the average spread around 500 nm. The autocorrelation plot in (E) shows the periodicity of 550-700 nm tangentially along the central axis of SYCP3. The estimated cluster diameter varied from 170-225 nm (see Sup. Fig. 2). The blue lines in the plot corresponds to 95 percent confidence bounds (+/- 0.08)

The general shape of chromatin is reproducible (Sup. Fig. 2) and presents characteristic clusters whose diameter is found to be in the range of 170-225 nm and periodicity to be in between 550-700 nm (see Fig. 2E, Sup. Fig. 2). In order to validate these clusters, we tested random distributions of chromatin (Sup. Fig. 3). The local condensation (clustering) in the SMLM image was characterised by calculating the distance to 20-500 nearest neighbours, and using the same method on the simulated data. The mean distance of 500 nearest neighbour was 86 nm for chromatin and 110 nm for the random data (Sup. Fig. 3), thereby indicating that the clustering seen in the real SMLM images of DNA arises from structural packing of DNA in the SC.

**Figure 3.**
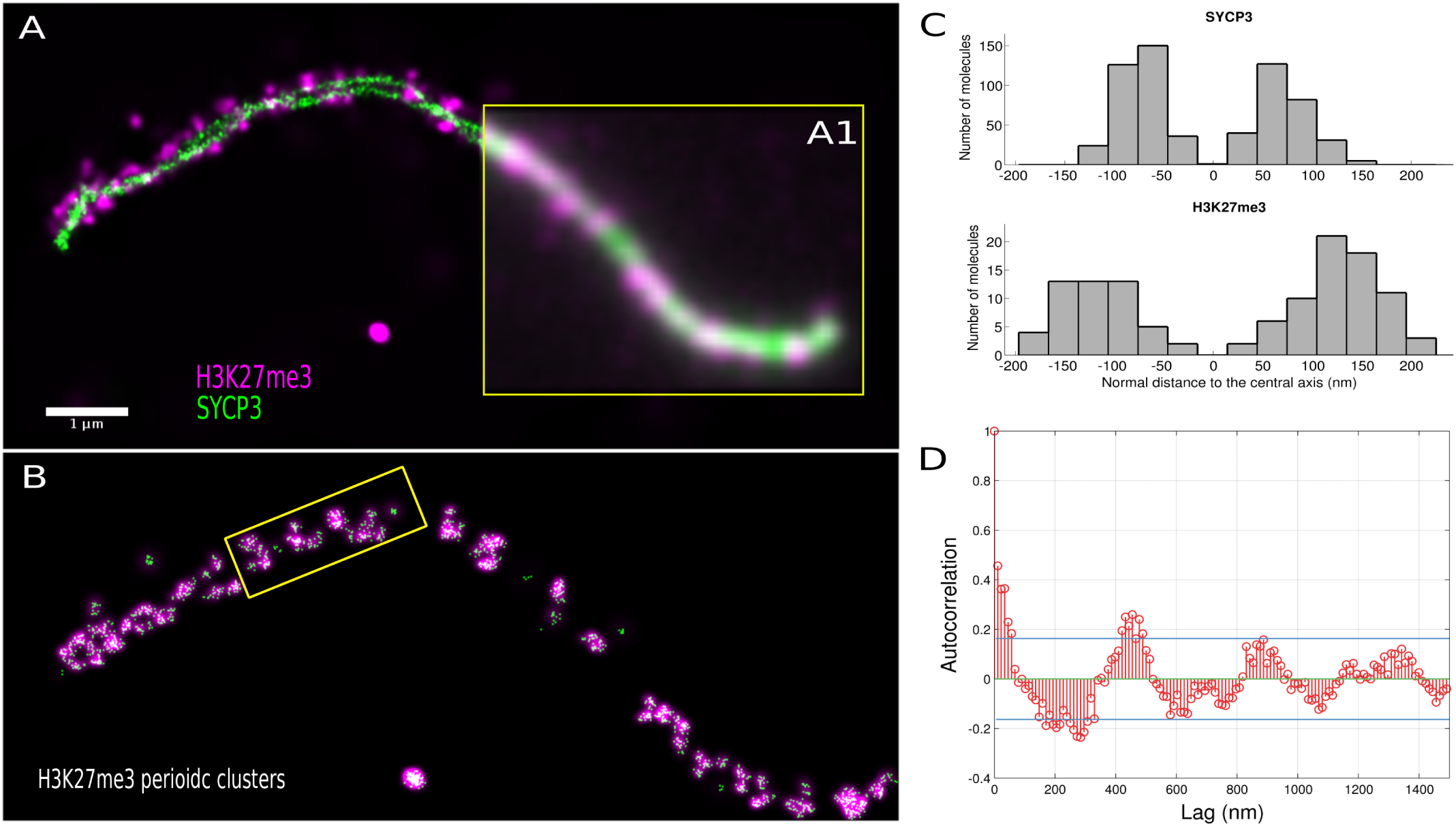
Periodic tangential clusters of transcriptionally inactive chromatin (H3K27me3): Two color SMLM image (A) immunostained with anti-SYCP3 (Alexa Fluor 555) and anti-trimethylated histone H3K27 (Alexa Fluor 488) with inset (A1) showing the widefield equivalent of the underlying localisation image. The white color results from superimposition of H3K27me3 and SYCP3 stainings. Large, periodic clusters of H3K27me3 mark associated with transcription repression, can be observed along the SYCP3 curve (B). In pink The clusters in (B) are shown in pink while in green are the single molecules belonging to the identified clusters The H3K27me3 clusters are found to be symmetrically distributed, laterally 50 nm apart from the SC, with a 90-130 nm diameter (C). The autocorrelation of the tangential distance of the clusters from the central axis of SYCP3 (D) indicates that the average periodicity of the cluster to be in range of ∼ 500 nm (Sup. Fig 4). The blue lines in the plot corresponds to 95 percent confidence bounds (+/- 0.1633)

### The repressive histone mark (H3K27me3) shows characteristic periodic clusters along the SC

Since histone modifications are known to regulate DNA participation in recombination and transcription, we tested if SMLM could be used to localize histone modifications in the context of chromatin. As was previously shown with confocal microscopy,^12^ the signals of repressive histone mark (H3K27me3) co-localized with SYCP3 at several points in our images as well. However, instead of colocalisation with SYCP3 over the whole length, we find large, periodic clusters (Fig. 3B) of H3K27me3 occurring at an average lateral distance of 40-50 nm from the strands of SYCP3, with occasional overlap (Fig. 3C). When comparing to previous results with confocal microscopy (and viewing our own widefield results), it is clear how colocalisation of SYCP3 and H3K27me3 would be assumed because of the ∼ 200 nm resolution of the imaging process. To estimate the periodicity and the size of the clusters, we computed an autocorrelation measure along tangential direction to the SYCP3. We observe that the clusters appear at an average distance of 450-650 nm (Fig. 3D, Sup. Fig 4), with the average cluster diameter between 90 to 130 nm (Sup. Fig 4). Staining of different chromosomes shows reproducibility of shape and hints at possible looping of chromatin around helical patterns of the SC (Sup. Fig 4). Given the H3K27me3 cluster spacing, it is possible that the H3K27me3 clusters may correspond to the chromatin clusters observed along the length of SC (Fig. 2C). Furthermore, due to the occasional symmetry of H3K27me3 clusters on both sides of SCs, we hypothesize that this histone mark might be associated with non-random sites along the genome.

### Centromeric histone mark (H3K9me3) labels one end of the SC

Mouse meiotic chromosomes are known to be telocentric, meaning that the centromeres are typically very close to one of the two telomeres of the chromosome, and these centromeric regions are associated H3K9me3. Seeing nanoscale spatial structure in H3K27me3, we next probed whether the large, condensed chromatin structures found at one end of the SC might be marked by H3K9me3, with unique nanoscale patterns. We co-immunostained SYCP3 and trimethylated histone H3K9 to study the localisation of the H3K9me3 mark with respect to SYCP3. Large, dense clusters of H3K9me3 could be seen at one of the ends of the pachytene chromosomes (Fig. 4), which we speculate to be the telocentric end. We hypothesize that the high density of H3K9me3 observed at the presumed telocentric end of SYCP3 (Fig. 4C) may be concomitant with the high chromatin density observed at the top axial end of the chromosomes (Fig. 2B), as the average axial spread of the large cluster in axial chromatin (Fig. 2B) and H3K9me3 (Fig. 4A) are both in range of 1-1.5 *μ*m. Moreover, we note that in some images, at the presumed non-telocentric end, the two SC strands seem to move apart, as observed previously.^23^ We have not observed such splitting at the presumed telocentric ends (Fig. 4, Sup. Fig. 5).

**Figure 4.**
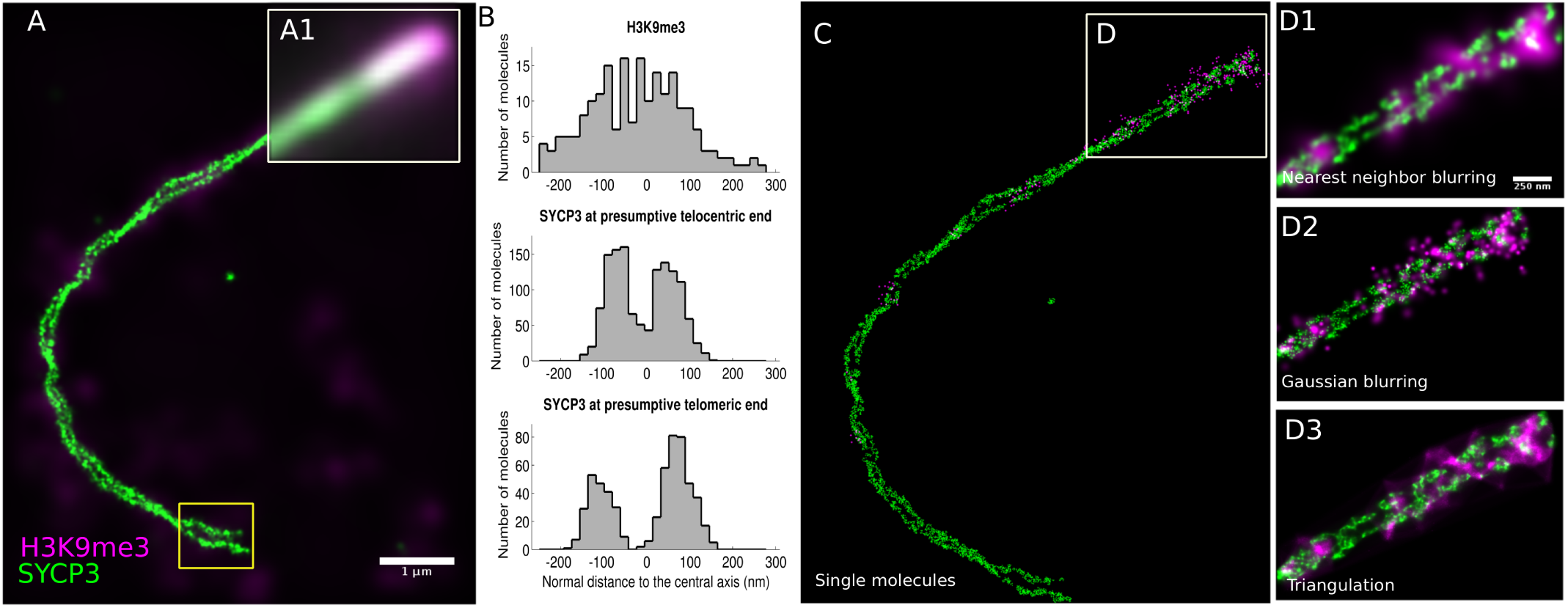
Telocentric distribution of H3K9me3 around SYCP3 molecules: Two color SMLM image (A) immunostained with anti-SYCP3 (Alexa Fluor 555) and anti-trimethylated histone H3K9 (Alexa Fluor 488) with inset (A1) showing underlying wide-field image. Large dense clusters of repressive centromere mark H3K9me3 can be seen at the presumptive telocentric end of the pachytene chromosomes. High H3K9me3 density at the telocentric ends of SYCP3 (C) is concomitant with high chromatin density at the top axial end in (Fig. 2B). (B) We observed that the strands of SYCP3 at the end where H3K9me3 co-localises (presumptive telocentric end) to be closer (∼ 135 nm) than at the presumptive telocentric end (∼ 180 nm) (Sup. Fig. 5). Point of representation of single molecules of SYCP3 and H3K9me3 is shown in (C). (D) compares different visualization strategies.

**Figure 5.**
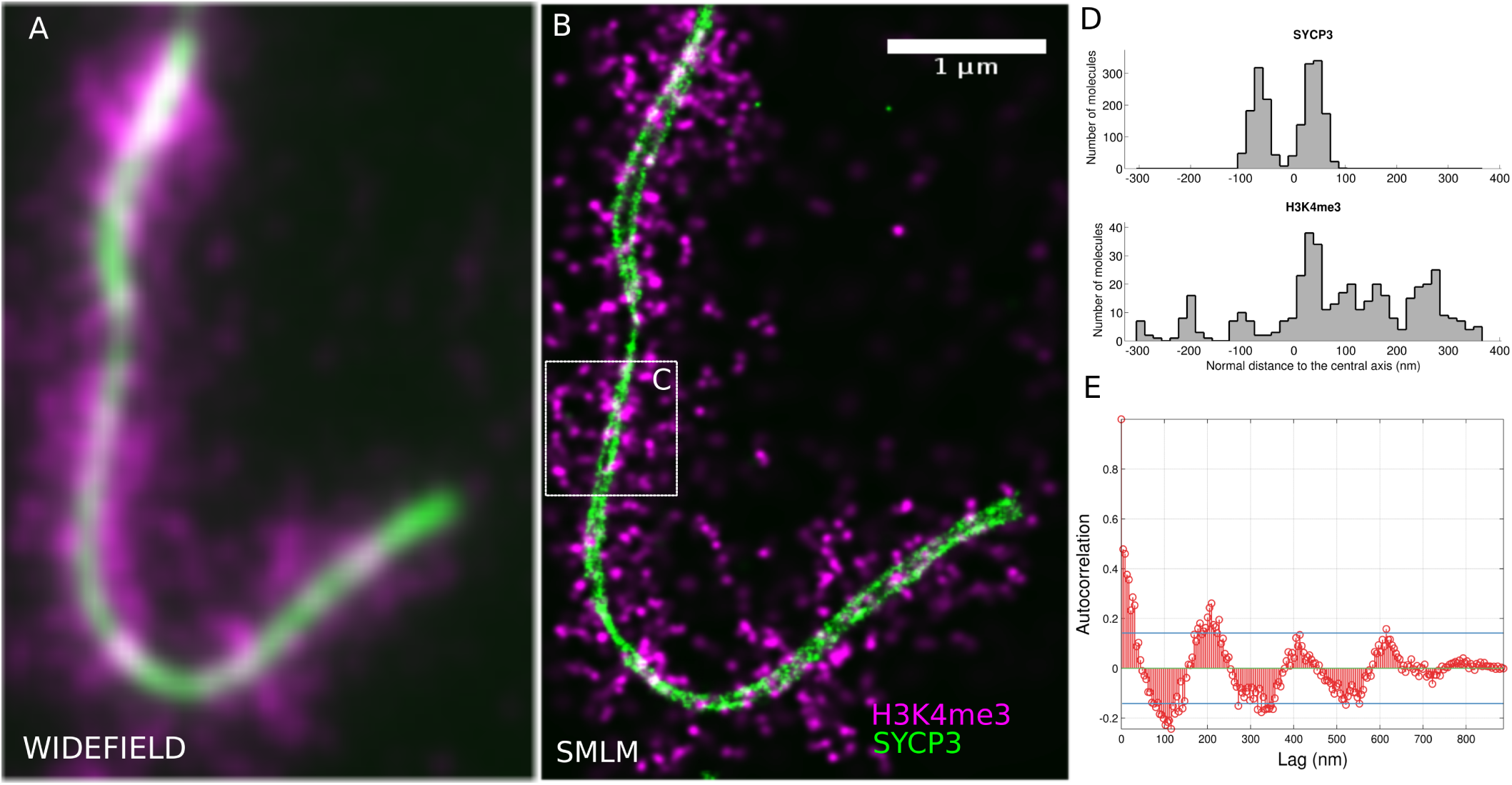
Radially emanating chromatin loops are punctuated with transcriptionally active histone mark (H3K4me3): Two color wide-field image (A) immunostained with anti-SYCP3 (Alexa Fluor 555) and anti-trimethylated histone H3K4 (Alexa Fluor 488) with (B) showing the corresponding SMLM image. shows radial loop like patterns emerging laterally to SYCP3. The distribution of H3K4me3 around SYCP3 is characteristically similar to chromatin distribution stained with Vybrant DyeCycle Violet (Fig. 2D). The average spread (∼ 300 nm) of radially emanating chromatin loops perpendicular to the lateral elements of SC is uneven across the SYCP3 strands punctuated with periodic patterns H3K4me3 mark (Sup. Fig. 6). Using autocorrelation function (D), we estimated the average spread of the radial loops to be larger than their spacing (∼ 200 nm) (Sup. Fig. 6).

### Histone mark (H3K4me3) associated with active transcription emanates radially from the axis of SC

Finally, we studied the distribution of H3K4me3, a histone modification associated with active transcription, along the meiotic chromosome. Surprisingly, we found the mark to be distributed both axially and radially, forming protrusions as long as 500 nm. Like H3K27me3, H3K4me3 appears discontinuous for the most part and tends to form tiny clusters (Fig. 5A-B, Sup. Fig. 6). The average spread of the radial emanations of H3K4me3 ranged from 300 nm to 500 nm with the overall distribution of H3K4me3 qualitatively similar to the chromatin distribution (Fig. 5D, Sup. Fig. 6, Fig. 2D). By autocorrelating the tangential distances from the central axis of SYCP3 (Fig. 5E), we estimated the average spread (∼ 500 nm) of the protrusions to be larger than their spacing (∼ 200 nm). Part of our images show a possible looping structure of H3K4me3 stained processes (Fig. 5C), compatible with data from electron microscopy of meiotic chromosome observed outside of mammalian class.^24^ Though chromatin extension have been observed before,^25^ to our knowledge, this is the first time that loops are observed in mammals especially in context of histone modifications.

**Figure 6.**
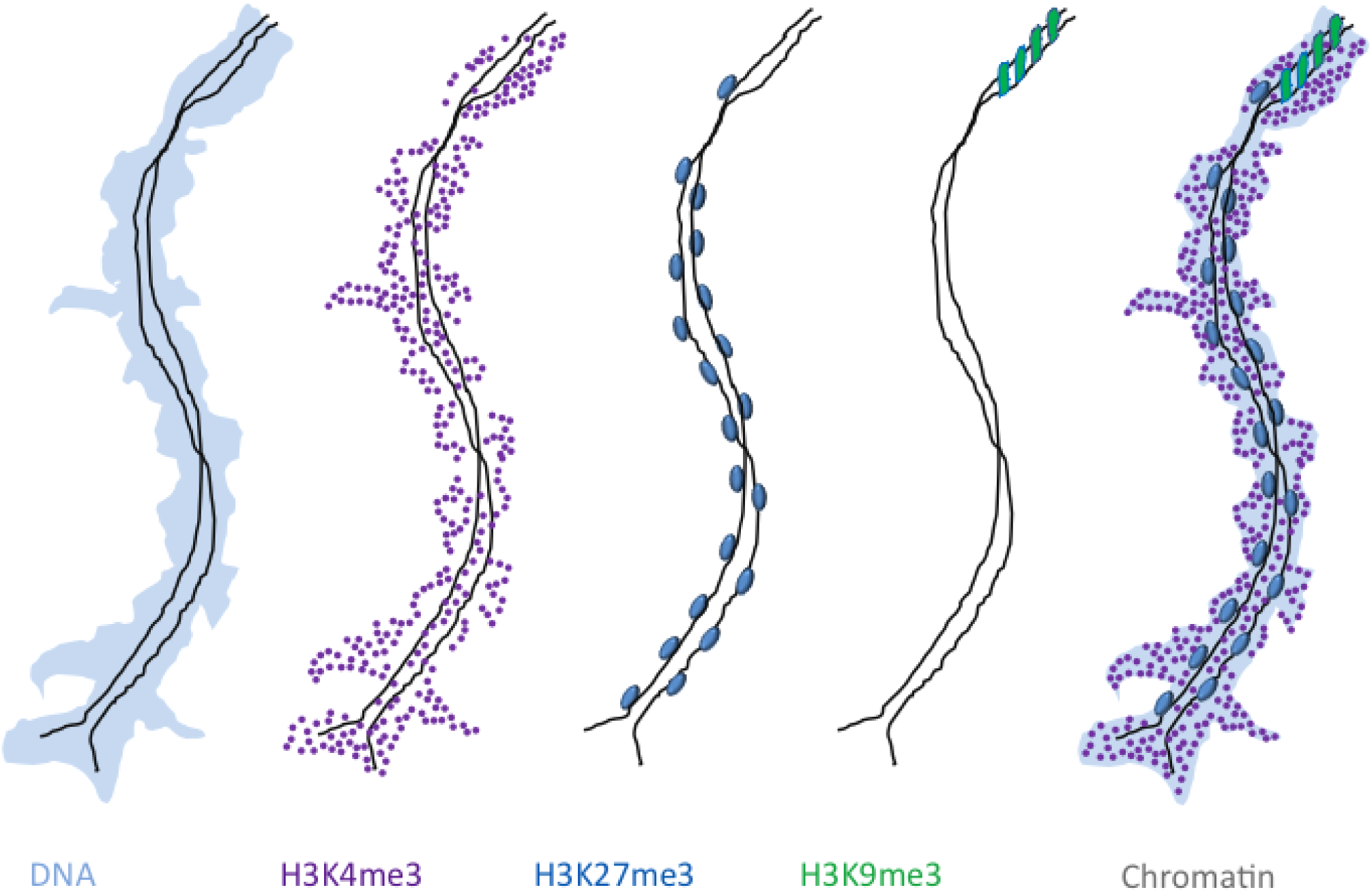
A model for spatial distribution of chromatin at pachytene stage of meiosis prophase I: Based on localisation maps of post-translational histone modifications, we can dissect the meiotic chromosome structure into at-least three distinct morphologies, highlighted by the differential, nanoscale organization: (1)Radial chromatin identified by trimethylation of histone H3 at lysine 4 (H3K4me3) – indicative of actively transcribed chromatin, (2) Tangential chromatin identified by trimethylation of histone H3 at lysine 27 (H3K27me3) – indicative of repressed chromatin and (3) Polar chromatin identified by trimethylation of histone H3 at lysine 9 (H3K9me3) – indicative of centromeric chromatin.

## 3 Discussion

We report the first single molecule resolution images of chromatin associated with pachytene chromosomes using light microscopy. Chromatin interestingly shows identifiable large clusters, that we speculatively associate with potential turns of the synaptonemal complex, modelled in other studies.^22, 26^ We also present the first high resolution images of histone modifications associated to the meiotic chromosome. Unexpectedly, histones show clusters possibly related to a level of chromatin organisation never reported before. Wide-field and SMLM images of H3K4me3 (Fig. 5A-B) show patterns that are morphologically compatible with characteristic loops associated with meiotic chromosomes. The function of these loops is still elusive, though one may speculate that they could participate to regulate recombination.^4^ H3K9me3 was found to localize at the presumptive centromeric regions. On closer examination, we note that some images hint at spiralization of H3K9me3-marked DNA at the extremity of synapsed chromosomes (Fig. 4D1-3).

Previously, it was reported that H3K27me3 is present exclusively at pachytene stage and co-localises with SC.^12^ We find periodic clusters of this histone mark near the axis of the synaptonemal complex. The pattern of H3K27me3 is especially interesting as it is occasionally symmetrical on each side of the SC, which hints towards an association with defined regions of the genome.^27^ These clusters may be selected to escape recombination, similarly to how DNA methylation was suggested to prevent recombination in transposon sequences.^28^ Another possible hypothesis is that H3K27me3 clusters may be involved in the formation of the SC itself.^29^

Our findings suggest a model for spatial distribution of chromatin at pachytene stage of meiosis prophase I (Fig. 6). Based on localisation maps of post-translational histone modifications, we can dissect the pachytene chromosome structure into three distinct spatial morphologies highlighted by the differential, nanoscale organization: (1) Radial chromatin identified by trimethylation of histone H3 at lysine 4 (H3K4me3) – indicative of actively transcribed chromatin, (2) Polar chromatin identified by trimethylation of histone H3 at lysine 9 (H3K9me3) – indicative of centromeric chromatin and (3) Tangential chromatin identified by trimethylation of histone H3 at lysine 27 (H3K27me3) – indicative of repressed chromatin. The overall view of the epigenetic make-up of the pachytene chromosome is consistent with increased chromatin accessibility away from the SC, and compaction of regions close to the SC. Relating this type of microscopy data to genomic data will be necessary to understand the biological function of the structural patterns we observe.

Our aim here was to demonstrate the power of single-molecule localization microscopy to describe the epigenetic landscape of pachytene chromosomes. Clearly, experimental evidence will be required to ascribe functions to the observed patterns of epigenetically marked chromosomal sub-domains. At a first step, it will be interesting to determine at which stage of prophase I the patterns of H3K4me3 and H3K27me3 are established, i.e. are they an exclusive feature of pachytene chromosomes or does a certain periodicity precede synapsis. By combining DNA, histone marks and markers of crossovers, it will be possible to localize crossover sites to epigenetic landmarks in single pachytene chromosomes. Beyond descriptive studies, the toolbox of genetic mouse mutants will help decipher how the radial and tangential chromatin domains are influenced by DSB formation and crossover resolution and vice versa. It is also conceivable to modify DNA in one locus and to observe the local consequences on chromatin structure, while the overall chromosome remains intact. Lastly, we cannot exclude that some of our observations are due to the spreading of pachytene chromosomes. Further investigations are needed to determine whether our findings hold true for in situ fixed pachytene cells. Since SMLM requires high laser powers resulting in fast photo-bleaching and labelling intensity is restricted, imaging of whole nuclei will require technical advances that go beyond the current set-up.

To conclude, SMLM imaging provides the first nanoscale localization map of three histone marks and chromatin on mouse pachytene chromosomes undergoing meiotic recombination. Observing histone modifications at the single molecule level will help define hypotheses for the functions of epigenetic modifications in meiotic recombination and in shaping higher-order genome architecture.

## 4 Methods and Material

The care and use of the mice were in accordance with the guidelines of the International guiding principles for biomedical research involving animals (CIOMS, the Council for International Organizations of Medical Sciences).

### Sample preparation of pachytene spreads

Pachytene spreads were prepared as described previosuly.^30^ Briefly, ovaries were dissected from E18.5 embryos and treated with hypotonic buffer (17 mM trisodium citrate-dihydrate, 50 mM sucrose, 5 mM EDTA, 0.5 mM DTT, 30 mM TRIS-HCl, pH 8.2) for 25 min. Ovaries were dispersed in 100mM Sucrose solution using 21G needles. The cell suspension was applied to poly-L-lysine covered slides and fixed in 1% PFA, 0.2% TritonX-100 solution, overnight in humidified chambers. Spreads were air dryed, washed in 0.1% TritonX-100 for 10 min, and three times in PBS. Slides were blocked in 5% BSA in PBS for 30 min at RT and stained with SYCP1 (ab15090), SYCP3 (ab97672) antibodies in 1:800 dilution at 37 degrees for 1.5 hours. For double-stainings of SYCP3 and histone-modifications, anti-H3K9me3 (ab8898), anti-H3K27me3 (Merck Millipore no. 07-449) and anti-H3K4me3 (ab8580) at concentrations of 1:1000 were used. Slides were washed three times 10 min in PBS and incubated with Alexa 488 and Alexa 555 secondary antibodies (LifeTechnologies), and washed again. Slides were embedded in ProLong Gold mounting reagent (LifeTechnologies). The DNA staining protocol has been described elsewhere.^17, 18^ Briefly, the slides were incubated with Hoechst 33258 (0.2 *μ*g/ml) for twenty minutes and washed with PBS briefly. In these cases slides were embedded in modified switching buffer consisting of glycerol with 10% imaging buffer (stock comprising 0.25mg/ml Glucose oxidase, 0.02mg/ml Catalase, 0.05 g/ml glucose in PBS). 10 *μ*l of the imaging buffer was placed on a glass slide after which the coverslip with the sample was placed upside-down on the buffer droplet. The coverslip was attached to the slide with either nailpolish or biologically inert dental paste (Picodent Twinsil) prior to imaging.

### Microscope setup

The configuration of the microscope set-up used here has recently been described.^17^ Briefly, for the present experiments, we used two diode-pumped solid-state (DPSS) lasers: 491 nm laser (Calypso 05 series, Cobolt, Sweden), 561 nm laser (Calypso 05 series, Cobolt, Sweden). The laser beam enters the microscope via mirrors and a collimator arrangement (5x expansion of beam diameter). The beam is focused by a lens into the back focal plane of an oil immersion objective lens (Leica Microsystems, 1.4 NA, 63x oil immersion with a refractive index of 1.518). The sample is actuated by a piezoelectrical stage for focussing. To achieve the high laser intensity for the localization mode, a lens is inserted in the optical pathway, leading to a smaller illuminated area (higher intensity) in the object region of interest, allowing for the appropriate conditions for the reversible photo bleaching. The emitted fluorescent light passes the dichroic mirror and is focused by a tube lens (1.0x, f=200mm) onto the charge-coupled device chip of a highly sensitive 12 bit black-and-white CCD chip (SensiCam QE, PCO Imaging, Kehlheim, Germany, with 102 nm effective pixel-size in the sample plane). A set of appropriate blocking filters (AlexaFluor 488 (Chroma, bandpass filter 525/50 nm), AlexaFluor 555 (Chroma, bandpass filter 609/54 nm) is mounted in a filter wheel in front of the CCD chip.

### Data acquisition and evaluation

For the dual color experiments, we sequentially imaged AlexaFluor 555 followed by AlexaFluor 488 (in case of H3K9me3, H3K27me3 and H3K4me3) or by Vybrant dye cycle Violet (in case of DNA staining). For imaging of AlexaFluor 555, 4000 frames with camera integration time of 90 ms were acquired with 90 mW of 561 nm excitation while imaging of AlexaFluor 488 was performed by acquiring 3000 frames with camera integration time of 100 ms at 90 mW with 491 nm excitation in the sample plane. The single fluorophore signal positions were determined by calculating the center of mass with a custom written software in Matlab (Mathworks).^31^ Temperature changes or mechanical relaxations can cause samples/instrument drift. A cross correlation algorithm was implemented to detect drift and extract shift vectors between reconstructed subsets (typically 100 frames). The extracted shift values were interpolated and used to correct the list of positions if an overall drift larger than 20 nm was detected. Measurements in dual-color microscopy usually suffer from chromatic aberrations. The chromatic shift between the two color channel images was extracted by reconstructing the individual color channels and calculating the cross-correlation between the reconstructions.

### Visualisation and data analysis

For visualization, we blurred each molecule position by a Gaussian distribution with a standard deviation corresponding to the mean value of the distance to the next 20 nearest neighbour molecule positions. In the generated image, the intensity corresponds to the local density of fluorophores resulting in an image equivalent to the one in conventional light microscopy.^32^ For high density single molecule reconstructions, structural resolution is dominated by localisation precision. Nearest neighbour visualisation is independent of localisation precision and helps in bringing out the underlying structure where the final reconstructions suffer from poor labelling density. A comparison with Gaussian blurring based on localisation precision, triangulation and point representation is shown in (Sup. Fig. 2). For quantitative data analysis, we converted the nearest neighbour blurred image to binary image based on global image threshold using Otsu’s method.^33^ We then extract all the single molecule positions underneath this binary mask for the downstream distance and autocorrelation analysis. Due to the curvilinear shape of SC, we first manually fitted central axis of the two strands with a polyline, segmentwise translated and rotated the coordinates of all the single molecules to get a more linear orientation. In order to get the inter-strand distance, we restricted the distance analysis to the sections where the two strands are clearly resolved and parallel to each other. Moreover, we rejected individual signals with distances larger than 500 nm (1000 nm for DNA) from the central axis to avoid false, unclear or background signals which did not belong to the complexes. The autocorrelation was calculated on tangential distances along the central axis. The formula for the autocorrelation of a spatial series *y*_*t*_ with a shifted copy of itself (shifted by lag *k*, where *k* = 0,…, *K*) in the case that *y*_*t*_ is a stochastic process is given by

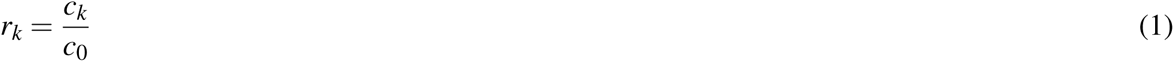

where *c* 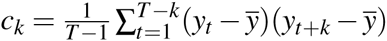 and *c*_0_ is the variance in the tangential distances.^34^ The blue lines in the autocorrelation plot correspond to 95 percent confidence bounds of the autocorrelation function assuming y is a moving average model.^34^

In our presentation, due to the unique spatial distribution of chromatin and its modifications, we used the Frenet frame to describe the geometric positions of the single molecules relative to the central axis of SC: (1) *lateral* to indicate single molecule positions that span the width of the SC, (2) *radial* to indicate positions of molecules that are perpendicular to the central axis of SC, and (3) *tangential* to indicate positions that are parallel to the central axis of the SC (for description refer to Sup. Fig. 7). We refrain from using the term axial other than in reference to the central axis of the SC because this term is already associated with description of the single strands of SC before synapsis.

## Acknowledgements

We thank Sapun Parekh for the help with the reagents, useful discussions and critically reading the manuscript. George Reid for the help with reagents and useful discussions. Wolf Gebhardt for the help with the reagents and Aleksander Szczurek for the help with the experiments. Work in K.T.K.’s laboratory is funded by the Austrian Academy of Sciences and by the European Research Council (ERC-StG-336460 ChromHeritance).

## Author contributions statement

K.P. conceived the project and planned the experiments. M.B. and K.T.K. prepared the samples. K.P. and S.R. performed the experiments. K.P. performed image data reconstruction and analysed the data. K.P. and G.B. wrote the software used for the data analysis. K.P., D.F., S.R., G.B., R.K., U.B. and C.C. analysed the results. K.P. and D.F. wrote the manuscript with input from all the other authors. D.F., K.T.K., R.K., and C.C. supervised the work. All authors read and approved the final manuscript.

## Additional information

The authors declare that the research was conducted in the absence of any commercial or financial relationships that could be construed as a potential conflict of interest.

